# Giant endogenous viral elements in the genome of the model protist *Euglena gracilis* reveal past interactions with giant viruses

**DOI:** 10.1101/2025.04.23.650285

**Authors:** Abdeali M. Jivaji, Sangita Karki, Emma Franken, Maria P. Erazo-Garcia, Zach K. Barth, Frank O. Aylward

## Abstract

Giant viruses in the phylum *Nucleocytoviricota* have increasingly been found integrated into the genomes of diverse eukaryotes. Here we report 8 Giant Endogenous Viral Elements (GEVEs) in the genome of the microalgae *Euglena gracilis*. The GEVEs bear signatures of genomic erosion, including invasion of transposable elements and duplications, suggesting that they are incapable of reactivation and virion production. Most of the GEVEs exhibit high average amino acid identity and cluster near each other in phylogenies of viral marker genes, suggesting that they are derived from the same initial viral lineage. Phylogenetic analysis of nucleocytovirus marker proteins reveals the viruses belong to the order *Asfuvirales* in the same broader lineage that includes African swine fever virus (ASFV), abalone asfarvirus (AbalV), and GEVEs recently found in the fungus *Rhizophagus irregularis* and the marine gastropod *Elysia marginata*, suggesting that widespread host range transitions have occurred in this lineage. This work expands the diversity of known endogenous giant viruses and expands the host range of the *Asfuvirales* to include the superkingdom *Discoba*.

## Introduction

Viruses in the phylum *Nucleocytoviricota* are dsDNA viruses colloquially called “giant viruses” due to many members of the phylum having particle sizes greater than 500 nm and genome lengths longer than 500 kb. Since the discovery of mimivirus in 2003, giant viruses have been found to be abundant and widespread in a variety of environments (1, 2). Many of the giant virus isolates have been characterized by infecting a few species of *Acanthamoeba* and *Vermamoeba* hosts (3, 4) which has significantly improved the molecular understanding of giant virus infection but belie the ecological impact of these interactions. Viruses such as Emiliana huxleyi Virus (EhV) (5), Ostreococcus tauri Virus (OtV) (6), Prymnesium kappa Virus (PkV) (7), and Paramecium bursaria Chlorella Virus (PBCV) (8–10) have been studied infecting the dominant organism in their respective environment and have provided insights into nutrient cycling and shaping microbial communities due to host-virus interactions (11, 12).

Moreover, the ecological impacts of giant viruses extend beyond microbial eukaryotes; viruses in the family *Iridoviridae* infect a diverse range of hosts, including protozoans, invertebrates such as insects, and even vertebrates such as fish and amphibians (13). Viruses in the orders *Chitovirales* and *Asfuvirales* are classical examples of giant viruses infecting vertebrates including mammals. The order *Asfuvirales* is exemplified by African Swine Fever Virus (ASFV) which is found to infect pigs (14). The amoeba-infecting Faustovirus and Pacmanvirus expanded the host range of *Asfuvirales* from the *Ophistokont* superkingdom to include *Amoebozoa* (15). The advent of next-generation sequencing and metagenomics has vastly accelerated the discovery of new giant viruses from a myriad of environments (2, 16–18). However, determination of hosts infected by giant viruses identified through metagenomics remains challenging and often relies on co-occurrence of the host and the virus, or recent gene transfers between viruses and their hosts (19).

Integration of viral genomes into those of their hosts has been widely studied in both prokaryotes and eukaryotes, and integration is often part of the natural infection cycle of the virus (20). Often, a virus will lose the ability to reactivate and become a genomic relic that gradually degrades in the host genome. The genes of the virus could potentially be co-opted by the host for other functions or merely degrade into “junk DNA” (21). A notable example of host co-opting viral genes is the syncytin protein in the placenta of mammals, which is derived from an ancient integration of a retrovirus (22). To date, the largest endogenous viruses discovered are derived from giant viruses, and they have been reported in a wide range of protist lineages. Giant virus genomes contain significantly more genetic material and novel genes for a host cell to co-opt once the integrated virus becomes degraded compared to smaller viruses. Moniruzzaman et al, (2020), found integration of giant viruses, termed as Giant Endogenous Viral Elements (GEVEs), in the genomes of diverse green algae (23), and later studies found additional GEVEs in still more chlorophyte genomes (24, 25). GEVEs have been reported in all superkingdoms of eukaryotes, and they are derived from multiple clades in the phylum *Nucleocytoviricota*. Sarre, et al, (2024), found evidence of 90 insertions of GEVEs belonging to medusavirus relatives in the genome of the holozoan *Amoebidium appalachense* (26). Denoeud, et al, (2024), discovered a large number of integrated *Phaeovirus* in the genomes of a wide range of brown algae (27). The record for the longest GEVE sequence is held by the genome of the mycorrhiza *Rhizophagus irregularis*, containing a 1.5 Mbp viral sequence belonging to the viral order *Asfuvirales* (28). Although in most cases it is unknown whether GEVEs are capable of reactivating to initiate a viral infection cycle and produce virions, a recent study provided evidence for a 617 Kbp long GEVE in the genome of *Chlamydomonas reinhardtii* (29). Since 2020, GEVEs have been detected in many microbial eukaryotes, expanding the list of known hosts to be infected by giant viruses because it is now generally assumed the presence GEVEs is due to a latent infection cycle.

Giant viruses have been shown to infect multiple superkingdoms of eukaryotes ranging from algae, amoeba, to vertebrates, yet giant viruses that infect some eukaryotic lineages remain unknown.

Particularly, the superkingdom of *Discoba* has just a single virus infecting a sole representative; the Bodo saltans Virus (BsV) in the order *Imitervirales*, which infects the marine kinetoplastid *Bodo saltans* in the phylum *Euglenozoa* (30). To expand the known hosts of giant viruses, we examined the possibility of finding a giant virus infecting the flagellate *Euglena gracilis;* it is a freshwater protist in the phylum *Euglenozoa*, class *Euglenida*, that is abundant in aquatic systems across the world. Euglena has also been the subject of study by biotech industries as a candidate for production of biodiesel and other important metabolites (31, 32). A high-quality genome sequence of *Euglena gracilis* had eluded researchers until Chen et al, (2024), published a chromosome-level assembly (NCBI Accession: GCA_039621445.1) (33).

Upon analyzing this assembly, we curated 8 viral regions that can be assigned as GEVEs. These GEVEs appear to have become degenerate as the markers seem to be absent or in multiple copies in many GEVEs. Phylogenetic analysis reveals the GEVEs belong to the order *Asfuvirales* expanding the host range of the order to include the superkingdom *Discoba*. Further research would be required to obtain an isolated giant virus infecting *Euglena gracilis* to understand the full extent of viral infection in the ecology of the freshwater protist.

## Results & Discussion

We detected 28 putative giant endogenous viral elements (GEVEs) in the genome of *Euglena gracilis* using ViralRecall 2.0 (34). We manually curated these to select for scores > 1 and remove regions composed of repetitive elements and false positives to ultimately arrive at a set of 8 viral regions that encode clear signatures of the phylum *Nucleocytoviricota*, such as homologs of the hallmark double jellyroll major capsid protein (MCP), A32 packaging ATPase, mRNA capping enzyme, multi-subunit RNA polymerase subunits, superfamily II helicase, and Virus Late Transcirption Factor 3 (VLTF3) (Table 1).

**Table 1:**
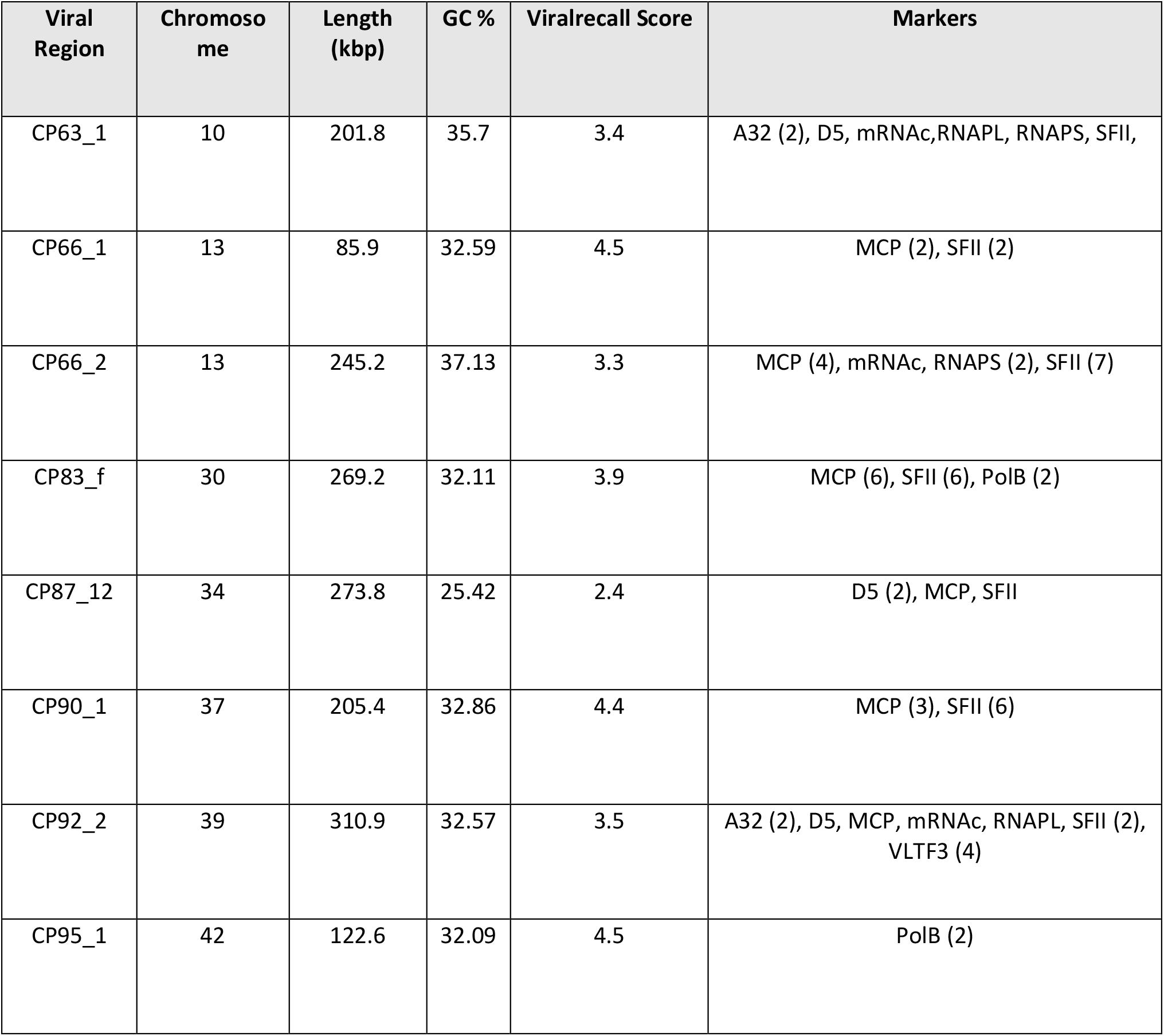
GEVE regions. A32: Packaging ATPase, mRNAc: mRNA capping enzyme, RNAPL/S: DNA-directed RNA polymerase alpha/beta subunit, MCP: Major Capsid Protein, SFII: DEAD/SNF2-like helicase, VLTF3: Poxvirus Late Transcription Factor VLTF3, PolB: DNA polymerase family B

| Viral Region | Chromosome | Length (kbp) | GC % | Viralrecall Score | Markers |
| --- | --- | --- | --- | --- | --- |
| CP63_1 | 10 | 201.8 | 35.7 | 3.4 | A32 (2), D5, mRNAC, RNAPL, RNAPS, SFII, |
| CP66_1 | 13 | 85.9 | 32.59 | 4.5 | MCP (2), SFII (2) |
| CP66_2 | 13 | 245.2 | 37.13 | 3.3 | MCP (4), mRNAC, RNAPS (2), SFII (7) |
| CP83_f | 30 | 269.2 | 32.11 | 3.9 | MCP (6), SFII (6), PolB (2) |
| CP87_12 | 34 | 273.8 | 25.42 | 2.4 | D5 (2), MCP, SFII |
| CP90_1 | 37 | 205.4 | 32.86 | 4.4 | MCP (3), SFII (6) |
| CP92_2 | 39 | 310.9 | 32.57 | 3.5 | A32 (2), D5, MCP, mRNAC, RNAPL, SFII (2),<br>VLTf3 (4) |
| CP95_1 | 42 | 122.6 | 32.09 | 4.5 | PolB (2) |

These viral regions range in length from 86-311 Kbp and collectively contribute ∼ 1.9 Mbp of genetic material to the genome. All the GEVEs exhibited a GC % content ranging from 25-35% which is significantly different from the *Euglena* genome GC% of ∼51% (Supplementary fig 1). We examined the tetranucleotide frequencies (TNF) of these regions compared to the adjoining sequence and confirmed that TNF signatures of the viral regions deviated significantly from the non-viral DNA in the same genomic loci, consistent with previous work that found distinct TNF signatures between host and GEVE sequences (Fig. 1A) (23). The GEVEs were also enriched for GVOGs than the neighboring DNA providing further confidence in our boundary delineation of GEVEs (Fig. 1A).

**Figure 1.**
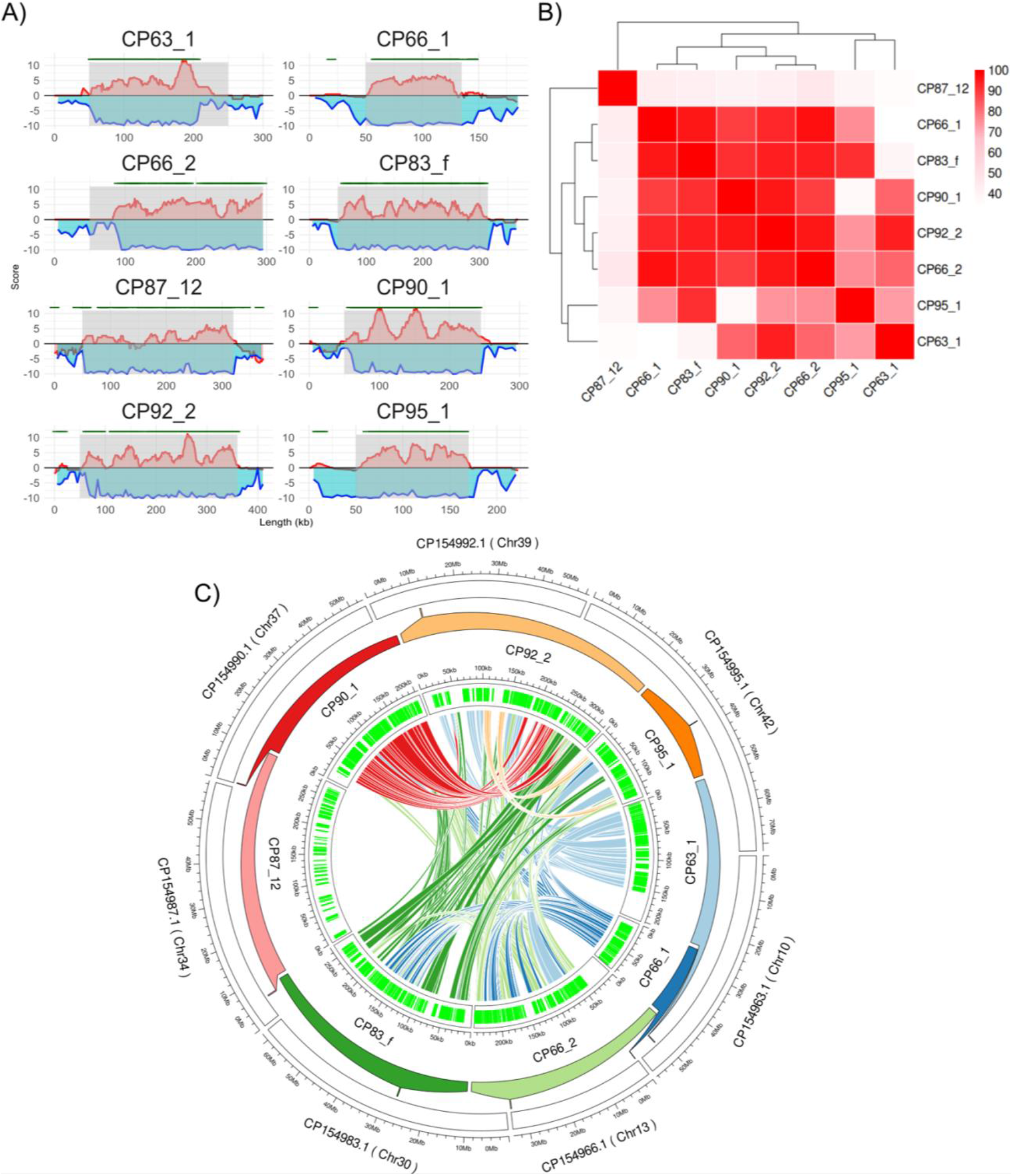
A) Viralrecall score (red), TNF deviation (blue) and GVOG positions (green) for the eight GEVEs (grey shaded) with 50 kb flanking region except CP66_2; B) All vs all Average Aminio acid Identity (AAI) matrix for the GEVEs. C) Circos plot showing GEVE (inner track) located on their respective *Euglena* chromosome (outer track). The lines represent a minimum 50 kb region of nucleotide similarity among corresponding GEVEs.

We compared the GEVEs to each other to understand their relatedness using Average Amino acid Identity (AAI) (35). The GEVEs form three clusters based on their AAI; CP87_12 is the most distinct GEVE with less than 50% identity with the rest of the viral regions, and GEVEs CP63_1 and CP95_1 form a separate cluster with some variation in the AAI % with other GEVEs. The rest of the GEVEs have relatively high pairwise AAI (> 70%), suggesting a recent shared evolutionary origin (Fig. 1B). The 8 GEVEs are situated on multiple chromosomes and seem to be nonspecific for any particular chromosomal loci. Interestingly, the viral regions CP66_1 and CP66_2 appear to be localized to the opposite ends of the same chromosome with CP66_2 forming part of the telomeric region (Fig. 1C). This is consistent with a study of GEVEs in Amoebidium, which found that they were spread throughout different chromosomes and regions within the same chromosome (26). The Euglena genome is also highly repetitive (>50%) and thus may be the target site for insertion leading to the dispersal of inserted GEVEs (33).

The coding density is high (>90%) for all GEVEs, consistent with the high coding density of free nucleocytoviruses. We looked at the presence of GVOGs and/or PFAM annotations for all predicted proteins. Approximately 50% of all proteins do not contain any hits in either database, and constitute a large pool of new proteins (Fig. 2 B).

**Fig 2:**
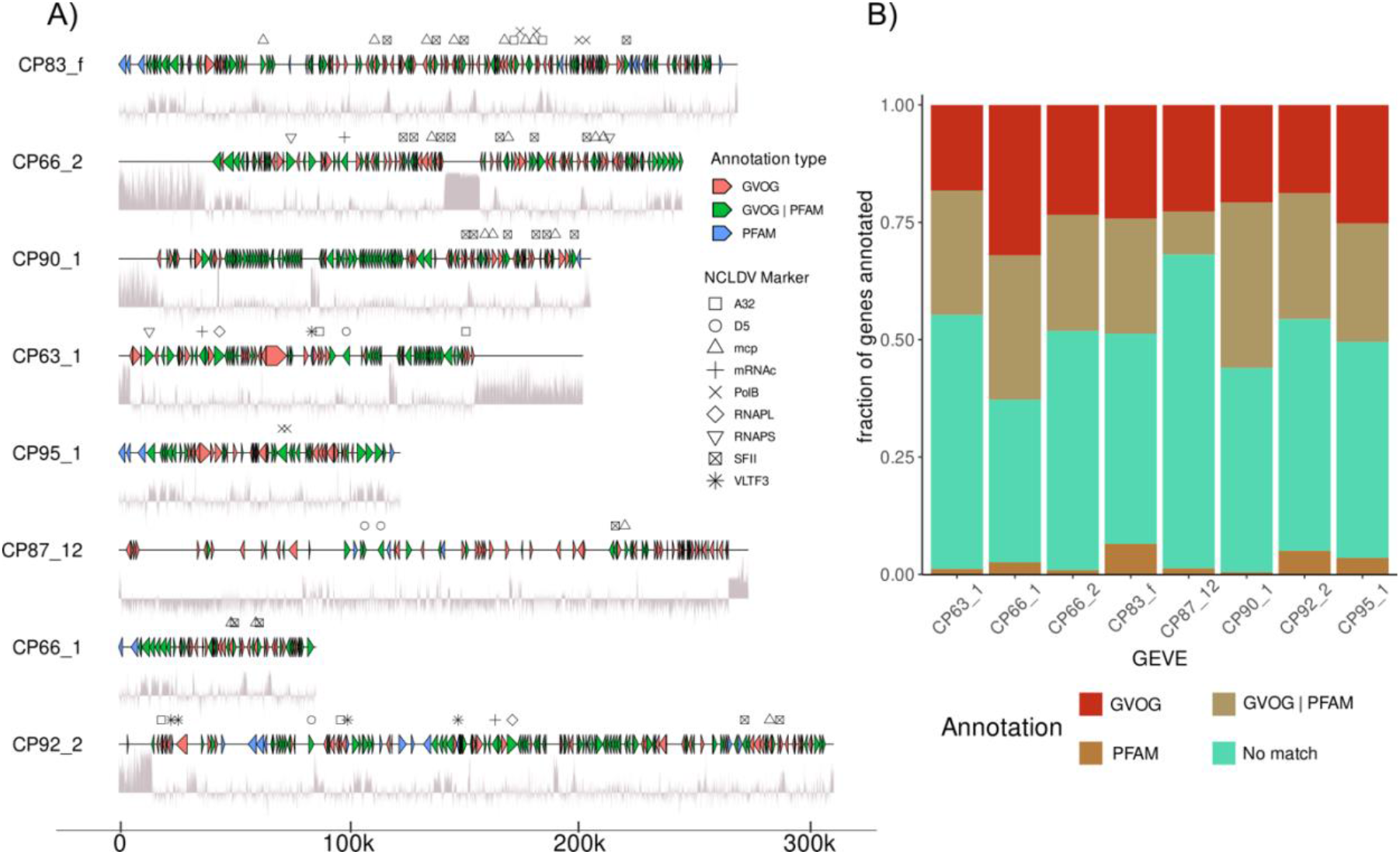
A) Gene maps of the 8 GEVE regions. The line under the GEVE represents GC% in a 50 bp sliding window. Only proteins with GVOG or Pfam annotations are shown. B) Fraction of genes annotated via GVOG or PFAM databases.

**Fig. 3.**
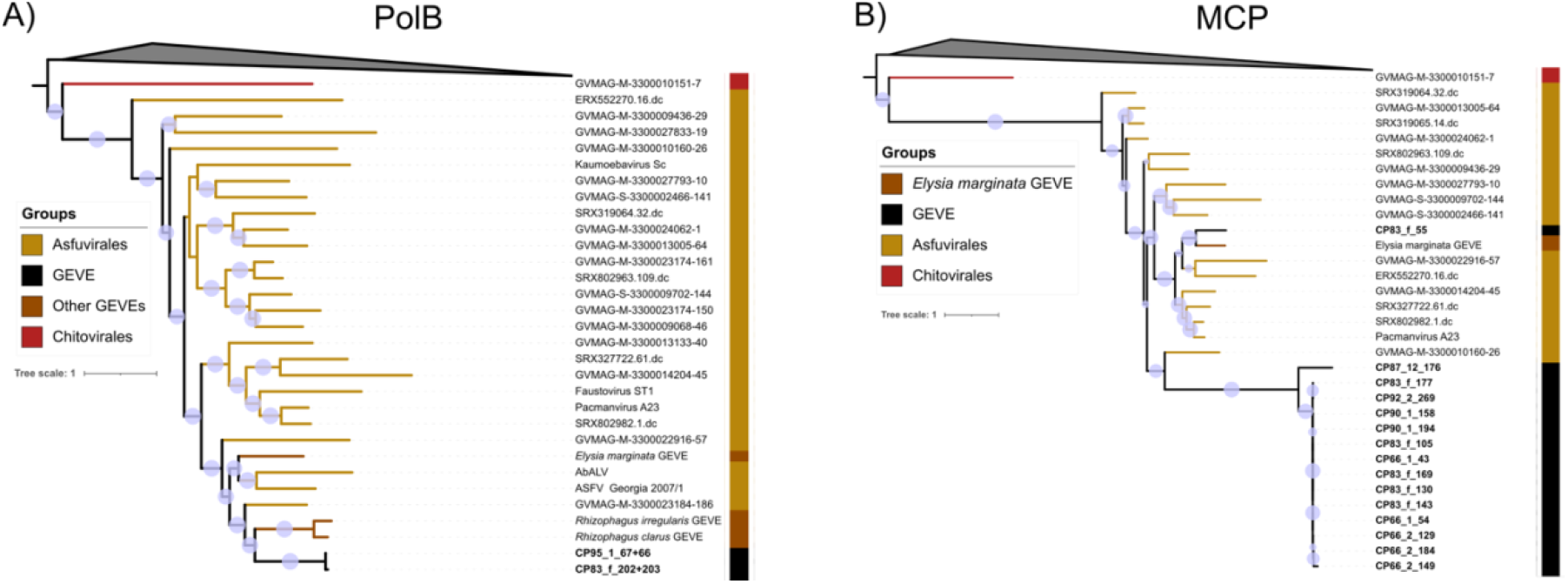
Phylogenetic trees of GEVE proteins PolB (A) and MCP (B) with respective proteins from all viruses in GVDB, and *Rhizophagus* and *Elysia marginata* GEVEs. The clades have been collapsed to only show the Order of giant viruses in which GEVEs clade. Bubbles indicate >90% bootstrap support.

### Gene synteny analysis and organization of genes and marker genes

Of the eight GEVEs, seven shared high amino acid identity (AAI, mostly >80%) indicating that they derive from the same viral lineage and potentially derive from multiple independent endogenization events.

These seven GEVEs also have a consistent %GC content of 32-35%, while the single outlier GEVE CP87_12) exhibited a lower GC% of 25%, further suggesting that this GEVE potentially derives from a distinct viral lineage. We also examined gene order within the GEVEs to gain insights into hypothesized shared origins for some of them. During our analysis we noticed that many of the markers on the GEVEs are present in duplicates. A common motif observed was the presence of MCP and SFII marker proteins arranged successively but transcribed in opposite directions. In the case of CP83_f, the gene clusters containing MCP and SFII along with additional neighboring genes are duplicated three times spanning the central 80 Kbp as can be seen in Supplementary fig. 2. These regions all share >95% nucleic acid identity, suggesting recent shared origins. Further, these motifs are spaced close to transposable elements that may act to mobilize these regions or serve as templates for recombination between GEVE regions. The abundance of repetitive elements and high similarity among the GEVEs has the potential to result into chromosomal rearrangements and introduce genetic variation, which has been observed in humans due to integration of Hepatitis B Virus (HBV) (36–38).

### Phylogenetic analysis

To examine the precise origins of the GEVEs, we performed phylogenetic analysis of these elements using various *Nucleocytoviricota* markers. Since the GEVEs exhibit signatures of genomic erosion, such as the presence of duplicate phylogenetic marker genes, a concatenated tree was not feasible. However, individual marker proteins in the Euglena GEVEs revealed that they fall within the order *Asfuvirales*. The Euglena GEVE markers fall within the same order of giant viruses in all phylogenetic trees constructed with high bootstrap support, providing further confidence in our conclusion. Recent work has reported GEVEs associated with the fungus *Rhizophagus irregularis* and the marine gastropod *Elysia marginata* (28), and we included these sequences in our trees as well.

Most GEVE MCPs are placed in the same branch, with the exception of that found in CP83_f_55, which is placed together with the MCP found in the *Elysia marginata* GEVE. These are within the same clade as the isolated Pacmanvirus that infects Acanthamoeba (39). The other GEVE MCPs, however, also seem closely related to Pacmanvirus MCP forming sister clades (39). Only CP83_f and CP95_1 contain PolB markers, and these form sister clades with PolB in the GEVEs in *Rhizophagus irregularis* and *Rhizophagus claris*. Surprisingly, the Euglena GEVE PolB sequences fall within the same clade with Abalone virus (AbalV), African swine fever virus (ASFV) indicating that the Euglena GEVEs belong to the family *Asfariviridae* along with ASFV and AbalV (41, 42). The clade also includes GEVEs found *Rhizophagus spp*. and *Elysia marginata*. Since PolB is an excellent marker used for taxonomic classification, these results demonstrate the very wide host range of viruses in the order *Asfuvirales*, which includes host*s* belonging to different superkingdoms of eukaryotes. Such rapid host-switching also complicates efforts to assess the deep evolution of nucleocytoviruses using their divergence times of their hosts as calibration points (43). For the same virus order to include viruses that infect such a wide range of hosts also magnifies the gap in research regarding the true host range of *Asfuvirales* (16).

### Functional Annotation and expression of viral proteins

We carried out the functional annotation of the GEVE proteins to determine the metabolic repertoire of the viruses infecting Euglena (Fig 4A). Most of the proteins did not contain hits in the KEGG and PFAM databases, although the GEVEs encoded many proteins involved in DNA replication and repair. The GEVEs CP63_1 and CP66_2 (CP63_1_8, CP66_2_60, CP66_2_193) contained proteins that had similarity to a nucleoid protein LMW5-AR found in African Swine Fever Virus (ASFV). The LMW5-AR protein encoded by A104R in ASFV is bacterial histone-like protein expressed in the late stage of viral replication and is involved in binding DNA and nucleoid formation and packaged in virion (44, 45). Multiple GEVEs (CP63_1, CP90_1, and CP92_2) also encode the Ulp1 protease which is a cysteine protease involved in processing of key polyproteins which form the core shell surrounding the nucleoid in ASFV. The ASFV protein (PS273R) is crucial during the late stage of viral replication and is packaged in the virion (44).

**Fig. 4.**
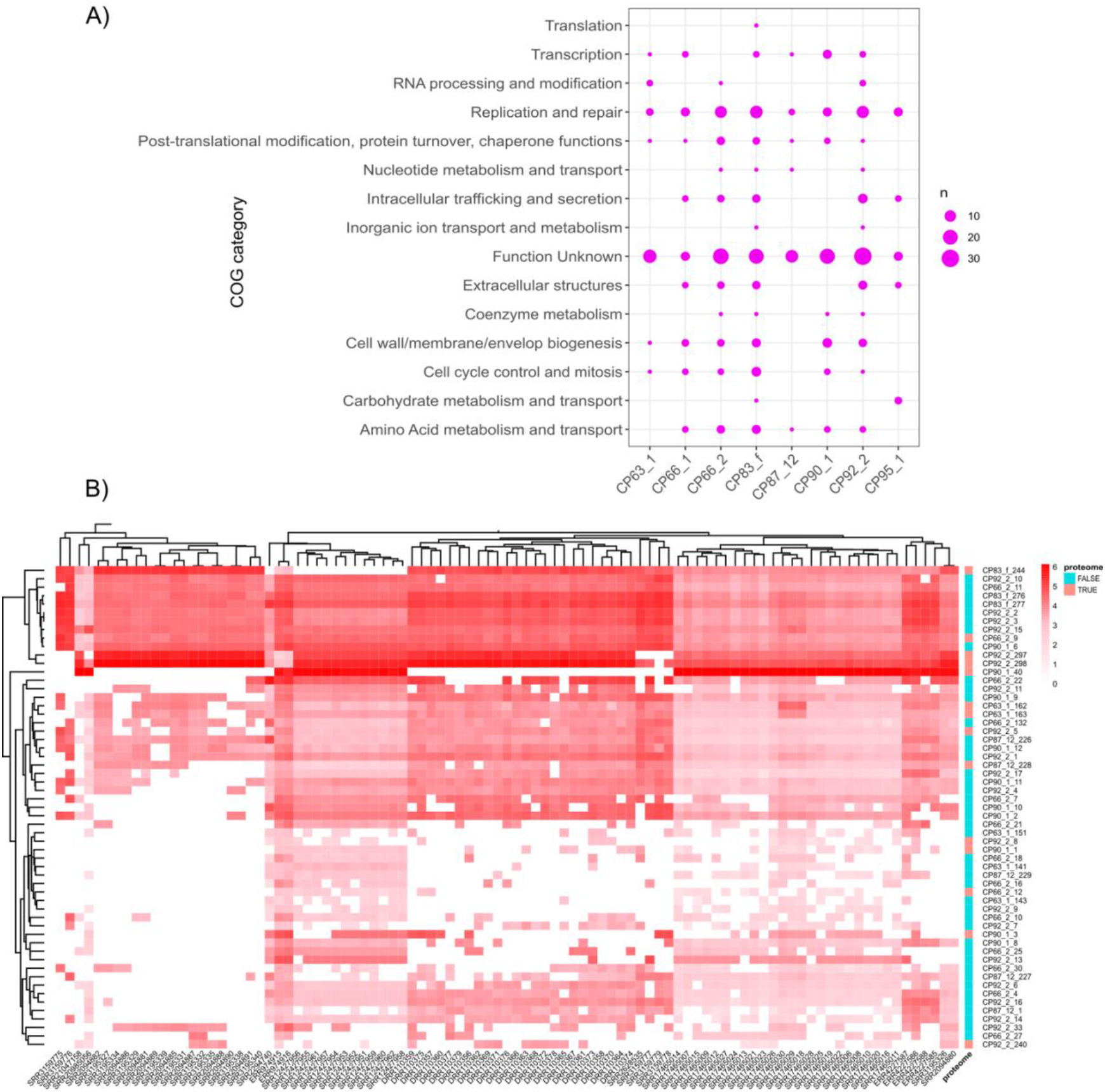
A) Functional annotation of GEVE proteins to show the COG categories of the GEVE proteins. B) Screening of 95 publicly available transcriptomes against all GEVE proteins. The heatmap has been subset to show proteins expressed in at least 20 out of 95 transcriptomes. The proteome column indicates GEVE proteins with at least one peptide hit in 5 publicly available proteomes of *Euglena gracilis*.

Out of the 315 annotated proteins, 33% match to Viruses and 88% of those viral proteins match to corresponding Asfarvirus proteins (Supplementary dataset). These key viral proteins involved in Asfarvirus replication found in the *Euglena* genome give us further confidence that the GEVEs are bona fide viruses in the family *Asfariviridae*.

To examine possible transcriptional activity of the GEVE genes, we screened the CDS of all predicted proteins in the 8 GEVE sequences against the 95 publicly available transcriptomes of *Euglena gracilis* in the SRA database on NCBI. Even if the GEVEs are unable to reactivate and produce virions, transcription of GEVE genes could indicate if any GEVE protein is being co-opted by the cell. To this end, we focused on transcripts found in a minimum of 20 out of 95 transcriptomes and considered these to be consistently expressed by the cell. Surprisingly, we found 57 genes in this set with 8 genes expressed in all 95 transcriptomes. HHPred analysis of these proteins reveal they belong to a diverse variety of functions (Supplementary dataset).

Many of these genes are predicted to encode for transposases/recombinase. On a deeper look, these recombinases were found to be distributed in abundance across the *Euglena* genome (supplementary fig 3), and those detected in GEVEs were largely restricted to the edges of the GEVE regions. Thus, these genes may represent selfish elements that have mobilized from the rest of the Euglena genome into either the GEVEs or regions immediately adjacent to them. We hypothesize that such a high abundance of transposons dispersed throughout the GEVEs might represent a defense mechanism of the cell that silences/degrades incoming foreign elements, though further work is needed to examine this avenue of research. Alternatively, many of the expressed integrases are present close to GEVE boundaries, and the virus may use the repeats at transposon insertion sites as integration sites to integrate the viral genome into the host DNA (46).

### Distribution of Euglena infecting virus in freshwater environments

We sought to determine if the viruses found as GEVEs in the *Euglena* genome can be found in the environment. To this end, we mapped short reads from 48 freshwater metagenomes. We selected the CP92_2 GEVE with 100 kb of flanking region as well as just the predicted GEVE region, as this is the most relatively complete viral region of the predicted GEVEs in Euglena. We decided to focus on freshwater metagenomes as *Euglena* is ubiquitous in freshwater aquatic systems.

We failed to find significant coverage of the GEVE (>20% covered fraction), with or without the flanking region, in any of the 48 metagenomes screened (47). This was not surprising as the GEVE, despite being relatively complete, still seems to be undergoing degradation in the genome. The genome is repeat-rich, and this is also observed in CP92_2, both tandem repeats as well as terminal inverted repeats. Despite the low coverage fraction over the whole length GEVE, we did observe significant read mapping to specific regions in the GEVE (Supplementary Fig. 4). These regions align with the location of TIRs indicating the reads mapped to repetitive or transposable elements. Thus, the repeats may be a part of some other organisms present in the environment or part of mobile genetic elements, but the read mapping to repeats may also happen by chance and thus it’s difficult to conclude anything significant.

### Methylation may play a role in silencing of GEVEs

We sequenced the *Euglena gracilis* UTEX 753 genomic DNA using nanopore to gain insights into methylation of GEVE regions. All the GEVEs had lower coverage than the surrounding DNA in our strain as observed in Fig. 5A. Despite lower coverage, the GEVE as well as the flanking region seems heavily methylated. The GEVEs, however, have a significantly lower GC% and contain a statistically significant higher fraction of methylated cytosines than the surrounding DNA.

**Figure 5.**
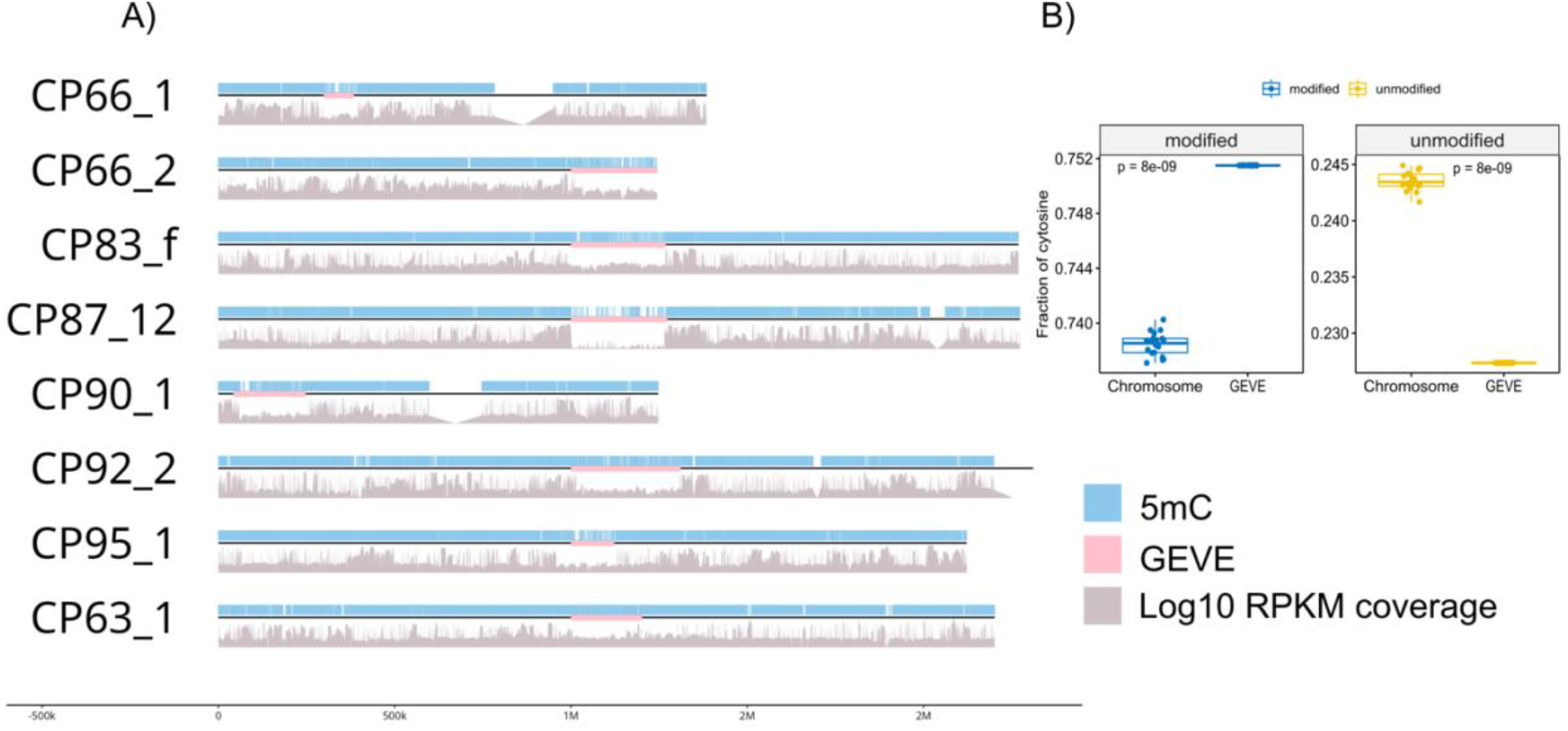
A) Cytosine methylation positions in the *Euglena* GEVEs (pink) with 1 Mbp flanking. B) Fraction of cytosine methylated and non-methylated on the Euglena chromosomes containing GEVE vs GEVE regions.

Methylation of cytosines is a crucial tool in the Euglena arsenal to fine tune responses to the environment. Even at low concentration of methylation modifying compounds, there is a drastic physiological response. This has been mainly investigated in proplastid regulation (48–51). Our analysis reveals that methylation of cytosine might also play a crucial role in antiviral response of *Euglena gracilis*. The GEVEs were also found to encode methyltransferases which might function as a defense mechanism against host silencing (26). An important piece of puzzle yet unexplored is the role of Base J, an unusual base modification found in euglenoids, and its utilization in silencing of GEVEs by kinetoplastids (52).

## Conclusions

The appreciation of endogenous giant viruses in eukaryotic genomes has increased dramatically with the increase in the number of high-quality eukaryotic genomes that have become available. Endogenous giant viruses, or GEVEs, have now been discovered in taxa spanning the majority of the superkingdoms of eukaryotes (23, 28, 29, 53). In the green algae *Chlamydomonas reinhardtii*, it was recently shown that a GEVE can reactivate and initiate a viral infection cycle, leading to the production of a small number of viral particles in an otherwise healthy culture (29). Although the characterization of GEVE reactivation in other systems will be needed to examine these dynamics further, it seems likely that the majority of GEVEs found in eukaryotic genomes are due to active integration of the virus as part of a cryptic infection cycle. Similar infection dynamics have also been found in mirusviruses associated with marine thraustrochytrids, and endogenous mirusviruses have been reported in a range of different hosts, suggesting that these cryptic infection cycles are common across both diverse protists hosts and a range of large DNA virus lineages (54–56). An analogy can be drawn between GEVEs and prophages found in the genomes of bacteria and archaea, which are widespread and play important roles in prokaryotic genome evolution. In most cases, the GEVEs appear to be silenced or degraded by the cell to prevent deleterious effects of viral activation. An example of silencing by cell is by methylating the GEVE regions as observed in Amoebidium (26). The GEVEs in this host were found to encode recombinases and chromatin modification enzymes such as DNA Methyltransferases (DNMTs) which might provide insights into endogenization as part of the viral life cycle or backup mechanisms to overcome silencing by the cell (26). Indeed, this also seems to be true for the GEVEs found in *Euglena gracilis*. Our analysis revealed they contain a significantly higher fraction of methylated cytosines than the surrounding DNA, thereby being silenced by host defenses. The viruses however also encode multiple methyltransferases which might act to overcome host silencing.

An alternative mechanism by which degradation of the GEVEs occurs is by either truncation, deletion, or invasion by mobile genetic elements and introns. Indeed, the first discovery of GEVE also noticed increased intron invasion into GEVE genes whereas the homologs in free virions were devoid of introns (23). Some of these introns are spliced appropriately and do not appear to impair viral reactivation, but others may be a signature of genomic erosion. Recently, McKeown, et al, (2025), identified seven *Phaeovirus* GEVE sequences in the freshwater brown alga *Porterinema fluviatile* and concluded that the GEVEs have undergone degradation due to truncation and elimination of core NCLDV marker genes; only one of the seven GEVEs contained all their demarcated core NCLDV genes. The GEVEs were also encroached upon by mobile genetic elements which lead to large scale disruptions and genomic rearrangements (53). All these strategies can also be observed to occur in the *Euglena* genome to silence the GEVEs. Once viral replication is quelled, integration of such a large chunk of DNA provides the recipient cell with enormous genetic potential ripe for co-option by the cell. However, given the abundance of ORFans and hypothetical proteins annotated in most giant viral genomes, elucidation of exact functions for GEVE proteins may require extensive biochemical investigation. GEVE proteins, thus, might open new avenues for eukaryotic genome evolution by providing seeds for novel pathways.

Finally, the family *Asfarviridae* in the order *Asfuvirales* has historically been limited to include only the isolate African Swine Fever Virus (ASFV). Matsuyama, et al, (2020), expanded the viruses in the family *Asfarviridae* with the discovery of Abalone virus (42). The discovery of Pacmanvirus and Faustovirus infecting amoeba added additional isolates to the elusive and understudied Order of *Asfuvirales* (15, 39, 57). Discovery of the 1.5 Mbp GEVE in the genome of *Rhizophagus irregularis* represents the longest genome of an Asfarvirus (28). The Hi-C data revealed that GEVE silencing by the cell is due to chromatin compaction of the GEVE region representing a novel mechanism of GEVE silencing. The detection of an Asfarvirus GEVE in the genome of the marine gastropod *Elysia marginata* further expanded the potential hosts infected by Asfarviruses (28). The PolB protein from Heterocapsa circularisquama DNA virus (HcDNAV) infecting the dinoflagellate *Heterocapsa circularisquama* contains high sequence similarity to the PolB of ASFV (58). This is a further expansion of the host-range of *Asfuvirales* Order to the *Alveolata* superkingdom of eukaryotes (59). With our detection of GEVEs in *Euglena*, this has the potential to expand the host range to include the superkingdom *Discoba* as potential hosts for the Order *Asfuvirales*. This exemplifies the broad-host range and extreme host-switching possible by these marvelous viruses. Their global distribution suggests that we might find these viruses playing a larger role in diverse ecosystems on Earth (16). The consensus distribution of all GEVEs in the *Euglena* genome belonging to the same order, *Asfuvirales*, leads us to conclude that cryptic viral infections are likely common in natural populations of *Euglena*.

## Methods

### Screening of Assembly

The *Euglena gracilis* assembly (NCBI Accession: GCA_039621445.1) was screened using ViralRecall 2.0 to discover GEVEs integrated into the genome (34). This resulted in 29 viral regions identified across multiple chromosomes. These were manually curated based on ViralRecall score (higher is better), minimum length of 50 Kbp, and number of viral hits assigned to each viral region. We also observed that three viral regions on chromosome 30 (CP154983.1), and two (out of the total three) viral regions on (CP154987.1) were adjacent to each other separated by a few hundred bp. These were joined manually to represent one viral region, respectively, to arrive at the final number of GEVEs in table 1.

### Average Amino Acid Identity (AAI)

Predicted proteins from GEVEs were compared to each other using custom scripts utilizing lastal to match proteins (60). The results were visualized using the R Pheatmap package (61).

### TNF Deviation and GC percentage

TNF deviation of the GEVEs was calculated in a sliding window of 5000 bp with custom R scripts using the R package BioStrings (62). GC percentage for the whole GEVE region was calculated using seqkit (63). For calculating GC percentage in a 50 bp sliding window, the seq-gc script from Thomas Hackl’s seq-scripts repository was used (https://github.com/thackl/seq-scripts).

### Synteny analysis

To examine the synteny of the GEVEs, the R package gggenomes was employed (64). The GVOG protein annotation was obtained from ViralRecall output. Briefly, the prodigal predicted ORFs were searched against GVOG HMM database using HMMER3 (65). The proteins were also matched against PFAM-A (release 37.0) database using eggNOG-mapper v2 using the parameters “--evalue 0.00001 --sensmode very-sensitive --pfam_realign denovo”, and independently using PfamScan (66, 67).

### Circular plot

To visualize the position of GEVEs with respect to the chromosome, we used the R package Circlize (68). The positions of GVOGs were obtained from ViralRecall output. To compare similar regions on the GEVEs, we used minimap2 and filtered out self-alignments and retained only alignments longer than 5 Kbp (69).

### Protein Homology search

To determine homology of each of the predicted proteins, we used a custom script using lastal to search against a custom database. The database combined RefSeq (Release 222), all giant viruses, and virophage/polinton proteins to increase the scope of the database. Analysis of the output was performed in R 4.4.1.

### Transcriptome search

To detect if any virus encoded protein was expressed, we downloaded 95 publicly available transcriptomes from NCBI using fastq-dump in SRA-tools (70). Coverm was used to map transcripts to the CDS of all ORFs predicted by prodigal, using the minimap2 mapper. Hits were reported in terms of RPKM. The values were converted to log10 RPKM to plot using R packages.

### HHPred search

The HH-Suite package was installed from its github repository (github.com/soedinglab/hh-suite). We also downloaded the UniRef30, PFAM, and PDB databases following instructions. All giant virus proteins were clustered using MMSeqs2 at 30% similarity and the output was used to create a custom database for use with the hhblits tools in HH-suite. Proteins were first matched against the giant virus protein database and the resulting alignments were used to search UniRef30, PFAM, and PDB databases by running hhblits with default parameters (71, 72). The script utilized GNU Parallel to increase efficiency (73). The .hhr files were tabulated using hhr2tsv script in Thomas Hackl’s seq-scripts repository (https://github.com/thackl/seq-scripts).

### Proteome search

Euglena proteomes were searched on the ProteomeXchange consortium, and the raw files were downloaded from FTP servers (74). Thermo .raw files were converted to indexed. mzML format using ThermoRawFileParser.exe (github.com/compomics/ThermoRawFileParser) with default parameters. The files were then searched against a protein database composed of the proteins obtained from transcriptome screening and the CRAP database of common contaminants, using FragPipe (75).

### Phylogenetic Analysis

For our phylogenetic analysis, we utilized Polymerase B, RNA polymerase large subunit, major capsid protein (MCP), and A32 marker genes. Marker sequences for the Euglena GEVEs were sourced from ViralRecall (34), while markers for the viral reference genomes were obtained using the ncldv_markersearch.py script (40). Fragmented Polymerase B sequences in Euglena GEVE were reassembled with the same script. We observed consistent hits to *Rhizophagus* spp. and *Elysia marginata* in the Euglena GEVE data; thus, we included these sequences based on BLAST results. We used Muscle5 (76) for sequence alignment and trimming was done using trimAl v1.4. rev15 with –gt 0.1 parameter (77). We constructed phylogenetic trees using IQ-TREE v2.1.2 (78) using LG+F+R10 model with the option -bb 1000 to generate 1,000 ultrafast bootstraps (79), -nt AUTO and --runs 5 to select the highest likelihood tree. All trees were visualized using ITOL (80).

### Genomic DNA extraction

*Euglena gracilis* Z (UTEX 753) was obtained from UTEX and grown in Euglena medium (UTEX) at 23°C in 16h:8h light:dark condition under shaking. 10^7 cells were harvested, and high-molecular weight genomic DNA was extracted using protocol from Gumińska et al 2018 (81).

### Nanopore sequencing and data analysis

Sequencing was performed in two rounds: Genomics Sequencing Center, Fralin Life Sciences Institute, Virginia Tech, using in-house library preparation and with Seqcoast Genomics. For in-house sequencing, library preparation was performed using SQK-LSK114 and DNA was loaded on a Promethion 114M flow cell. The pod5 files from both sequencing runs were pooled and basecalling was performed using Dorado v0.9.1 basecaller using parameters –min-qscore 9, and models hac 5mCG_5hmCG, 6mA models for modified basecalling against the *Euglena gracilis* reference genome assembly (See Data Availability). The modified bases were tabulated using modkit pileup v0.3.2 and summarized using modkit summary. Data analysis was performed in R using packages gggenomes and ggpubr.

## Acknowledgements

One round of sequencing was performed at the Genomics Sequencing Center which is part of the Fralin Life Science Institute at Virginia Tech. We thank Rich Helm for assistance with the proteomics data analysis. We acknowledge the use of the Virginia Tech Advanced Research Computing Center (VT ARC) for bioinformatic analyses performed in this study Funding: National Science Foundation CAREER award no. 2141862 (F.O.A.), National Institutes of Health R35 grant no. 1R35GM147290-01 (F.O.A.).

## Data Availability

The supplementary figures, supplementary dataset, and the bam file for the methylated basecalling has been deposited in Zenodo archive, doi: 10.5281/zenodo.15238724.

